# Interdomain disulfide bonds of rabbit kappa light chain allotypes influence mouse-rabbit chimeric antibody performance

**DOI:** 10.1101/2025.09.17.676883

**Authors:** Yang-Nim Park, Douglas W. Houston

## Abstract

Chimeric monoclonal antibodies have emerged as powerful tools to extend the capabilities of traditional monoclonal antibodies. These antibodies are made by replacing the variable region of a monoclonal antibody with the variable region of another antibody, typically from a different species, enabling use in a wider range of applications. Although theoretically compatible, interspecies differences in antibody structure can complicate chimeric antibody design and performance. In this study, we evaluated the impact of rabbit light chain allotype on the expression and function of chimeric antibodies containing mouse light chain variable regions. We found that constructs using the rabbit kappa 1 b4 (K1-b4) allotype frequently exhibited poor recombinant expression and, in some cases, lost antigen recognition. Structural analysis implicated disruption of an intrachain, interdomain disulfide bond as a contributing factor. Restoration of key residues predicted to re-establish this bond partially rescued both expression and activity. Additionally, chimeric antibodies incorporating the rabbit kappa 1 b9 (K1-b9) allotype, which contains a disulfide bond not disrupted by mouse variable region sequences, consistently maintained robust production and antigen-binding activity across multiple applications, including immunoblotting, immunoprecipitation, and immunostaining, for several test antibodies. Our findings underscore the importance of light chain scaffold selection in recombinant antibody engineering and provide practical guidance for optimizing chimeric antibody design to preserve both expression and function.

## Introduction

The “exquisite specificity” of monoclonal antibodies (mAbs) toward their antigens has made these molecules invaluable for biomedical research, diagnostics, and for developing therapeutic drugs for human diseases, including cancers, autoimmune disorders, and infectious diseases (1–3). Monoclonal antibodies are commonly generated using hybridoma technology, in which antibody-producing B-cells from immunized animals are fused with compatible immortal myeloma cells (4). This approach has revolutionized the production of monoclonal antibodies of defined specificity on a large scale.

However, there are limitations associated with this well-established technology. Notably, the long-term stability of hybridomas requires cryostorage and careful maintenance efforts to mitigate loss of production (5). Also, hybridoma technology is primarily applicable to rodents, owing to a general lack of fusion-compatible myeloma cells for other animal species. An emerging trend is the generation of recombinant antibodies, obtained either by cloning the hybridoma immunoglobin loci (or by cloning directly from B-cells), or through in vitro screening of libraries of antibody variable regions (6–8).

Additionally, the utility of recombinant antibodies can be increased through the generation of chimeric monoclonal antibodies. These molecules combine the antigen specificity of an antibody generated in one species with the effector functions of another, enabling broader applications in research, diagnostics, and therapeutics. Notably, recombinant monoclonal antibodies were first historically made as chimeric mAbs, for which variable regions of a mouse monoclonal antibody were fused with the constant regions of a human IgG molecule (9), resulting in a humanized antibody. More advanced variants followed using complementary-determining region (CDR) grafting (10), which involves more precise replacement of the human IgG CDRs with those of a mouse monoclonal antibody.In general, reducing the amount of ‘foreign’ constant domain sequences in the chimera would serve to minimize the immune response when used in a patient, a prerequisite for effective antibody-based drug development, but also helps eliminate inappropriate detection by secondary antibodies against the ‘donor’ species of the variable regions.

Recent advancements in genomics technologies have enabled large-scale antibody sequencing and subsequent recombinant and chimeric antibody creation, facilitating diverse antibody engineering (11). These molecules can be generated in various formats, including single-chain variable fragments (scFv), single-chain variable Fc-containing (scFc), and antigen-binding fragments (Fab), as well as intact original and chimeric antibodies (2; 11; 12).

Whereas these techniques are widely used in the development of human therapeutics, recombinant chimeric mAbs are increasingly being made for research purposes, using chimeras between mouse and rabbit, or involving other species. These chimeric approaches not only enhance the use of antibodies in research, allowing multiplex detection of antigens using defined and renewable mAb reagents, but also generate insights into sequence constraints for mAb specificity and stability.

Recombinant chimeric mAbs made using rabbit antibody scaffolds pose some challenges, however. In contrast to multiple heavy chain isoforms found in mice and humans, the rabbit immune repertoire contains primarily one heavy chain (13). Also, whereas mice and humans have two light chain isotypes — the kappa (*κ*) isoform predominates over lambda at ratios 95:5 and 60:40 in mice and in humans, respectively — rabbits have two kappa isoforms (*κ* 1 and *κ* 2) and one lambda isoform. Furthermore, the kappa 1 isoform (K1), comprising 70-90% of rabbit light chains, has four allotypes (b4, b5, b6, and b9; **Table 1**) (14; 15), that exhibit distinct patterns of intrachain, interdomain disulfide bonds (16). These bonds are thought to give increased stability to these rabbit isoforms (17–19), but are absent in rabbit kappa 2 and lambda, as well as in mouse and human light chains. The K1-b4,-b5, and -b6 allotypes exhibit disulfide bridges between Cys80 and Cys171, whereas in the K1-b9 allotype, these bridges instead exist between Cys108 and Cys171 (16).

**Table 1:** Light chain isotypes for mouse, human, and rabbit IgGs.

|  | Mouse and human |  | Rabbit |  |  |  |  |  |
| --- | --- | --- | --- | --- | --- | --- | --- | --- |
|  | Kappa | Lambda | Kappa1 | Kappa1 | Kappa1 | Kappa1 | Kappa2 | Lambda |
| Allotype | N/A | N/A | b4 | b5 | b6 | b9 | N/A | N/A |
| Number of interdomain disulfide bond (idSS) | 0 | 0 | 1 | 1 | 1 | 1 | 0 | 0 |
| Amino acid positions of idSS cysteines | N/A | N/A | 80 and 171 | 80 and 171 | 80 and 171 | 108 and 171 | N/A | N/A |

*The rabbit kappa 1 light chain allotypes comprise 70-90% of rabbit light chains and contain interdomain disulfide bonds not found in other rabbit light chains or other species (14; 15).*

## Results

### Generation of chimeric recombinant antibodies

To begin to archive and curate the sequences of unique hybridomas in the DSHB collection and generate cognate recombinant and chimeric antibodies, we obtained permission from the original depositor and sequenced the VL and VH domains from two DSHB hybridomas producing mAbs against transcription factors PAX6 and PAX7 from chicken (20) (anti-PAX6 [PAX6], anti-PAX7 [PAX7]). These antibodies recognize the target proteins across a range of vertebrate species, are effective in many different applications, including immunoblotting, immunoprecipitation, and immunofluorescence, and have been shown to be specific using knockout samples (e.g., (21; 22)). We also compared these chimeric recombinant antibodies with recombinant antibodies made against Hemagglutinin (anti-HA [12CA5], from Influenza A virus strain A/Victoria/3/1975 H3N2; (23)) and against GFP, using a mAb developed by DSHB (anti-GFP [DSHB-12E6]; (24)).

To generate recombinant mAbs, the endogenous signal peptides were retained and the variable light (VL) and variable heavy (VH) domains were each cloned into vectors containing the mouse kappa *(κ*) light chain and mouse IgG1 heavy chain, respectively, to enable expression of recombinant mouse IgG1 (**Fig. 1A**). To generate mouse-rabbit chimeric recombinant antibodies (**Fig. 1B**), the same VL and VH domains were fused to rabbit kappa light chains (K1-b4 allotype; K1-b9 allotype) and to the rabbit IgG gamma heavy chain (the only isotype in rabbits), respectively.

**Figure 1:**
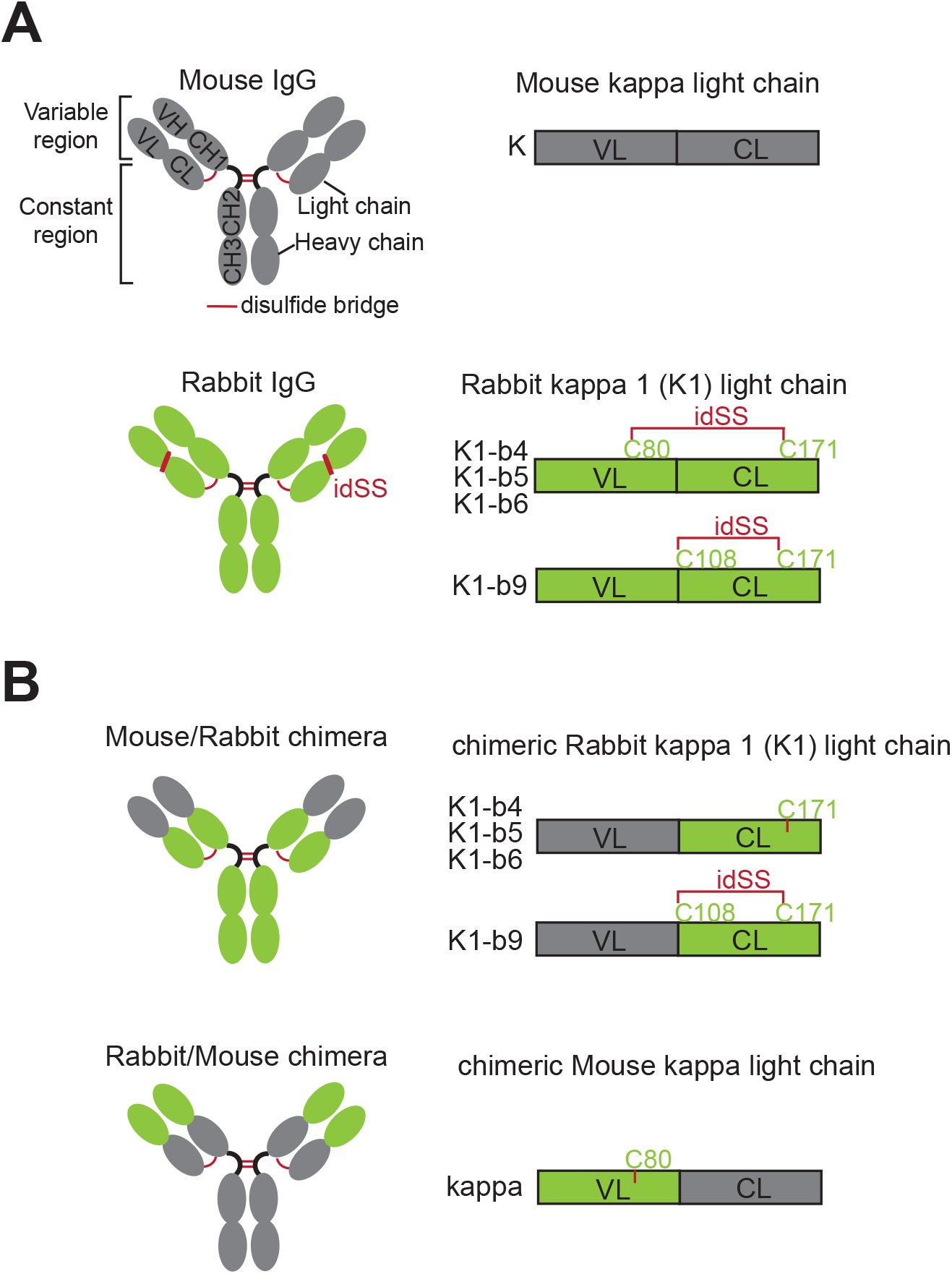
Structure and generation of recombinant and chimeric antibodies. (A) Schematic depiction of mouse IgG1 antibody high-lighted with the presence of disulfide bridges (red). Positions of relevant cysteines and interdomain disulfide bonds (idSS) between rabbit kappa light chain variable (VL) and constant (CL) regions are indicated. (B) Chimeric recombinant mouse/rabbit IgG or rabbit/mouse antibodies are generated by fusing variable regions with constant IgG regions of the desired species.

To assess recombinant and chimeric recombinant antibody production, heavy and light chain-containing vectors were co-transfected (1:2 ratio, heavy chain:light chain) into suspension cultures of two strains of HEK293 cells (Expi293F and HEK293C18). Supernatants were harvested after incubation for seven days, and the production of antibodies in the supernatants were quantified using ELISA. The activity of the antibodies was then further assessed (see subsequently). The entire process of generating recombinant antibodies in this study is illustrated stepwise in **Fig. S1**. Importantly, control experiments showed that when used in combination with anti-mouse antibodies, chimeric rabbit antibodies containing mouse variable regions were not recognized by anti-mouse secondary antibodies (**Fig. S2**).

In HEK293C18 cell suspension cultures, the average (mean) yields of anti-HA, anti-GFP, anti-PAX6 and anti-PAX7 antibodies in the supernatants ranged from 6.0 - 87.0 µg/ml (**Fig. 2A**; **Table 2**). In contrast, the production of the corresponding chimeric recombinant antibodies was markedly lower, ranging from 0.3 - 1.4 µg/ml (**Fig. 2A**; **Table 2**). To assess the extent that these reduced production levels might result from instability or altered processing of the chimeric recombinant immunoglobins, we first examined their relative migration in reducing and non-reducing SDS-PAGE gels. Under both reducing and non-reducing conditions, the recombinant mouse mAbs displayed banding patterns similar to native hybridoma immunoglobulin, with bands of approximately 50 kilodaltons (kDa) and 25 kDa (relative migration), corresponding to the intact heavy and light chains in reducing gels (**Fig. 2B**), and with normal, complete forms of the antibodies between 120 and 150 kDa in non-reducing gels (**Fig.2B**). However, the mouse-rabbit chimeric recombinant antibodies showed weaker and less distinct bands under both conditions, with broader and slightly slower migrating bands on non-reducing gels (**Fig. 2C**).

**Table 2:**
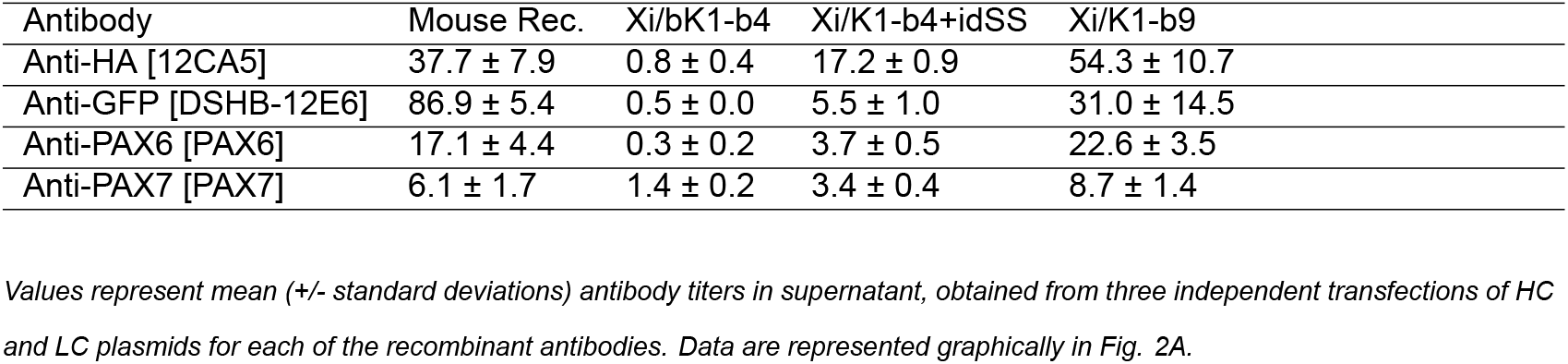
Expression levels of recombinant and chimeric recombinant antibodies in HEK293C18 cell suspension cultures.

**Figure 2:**
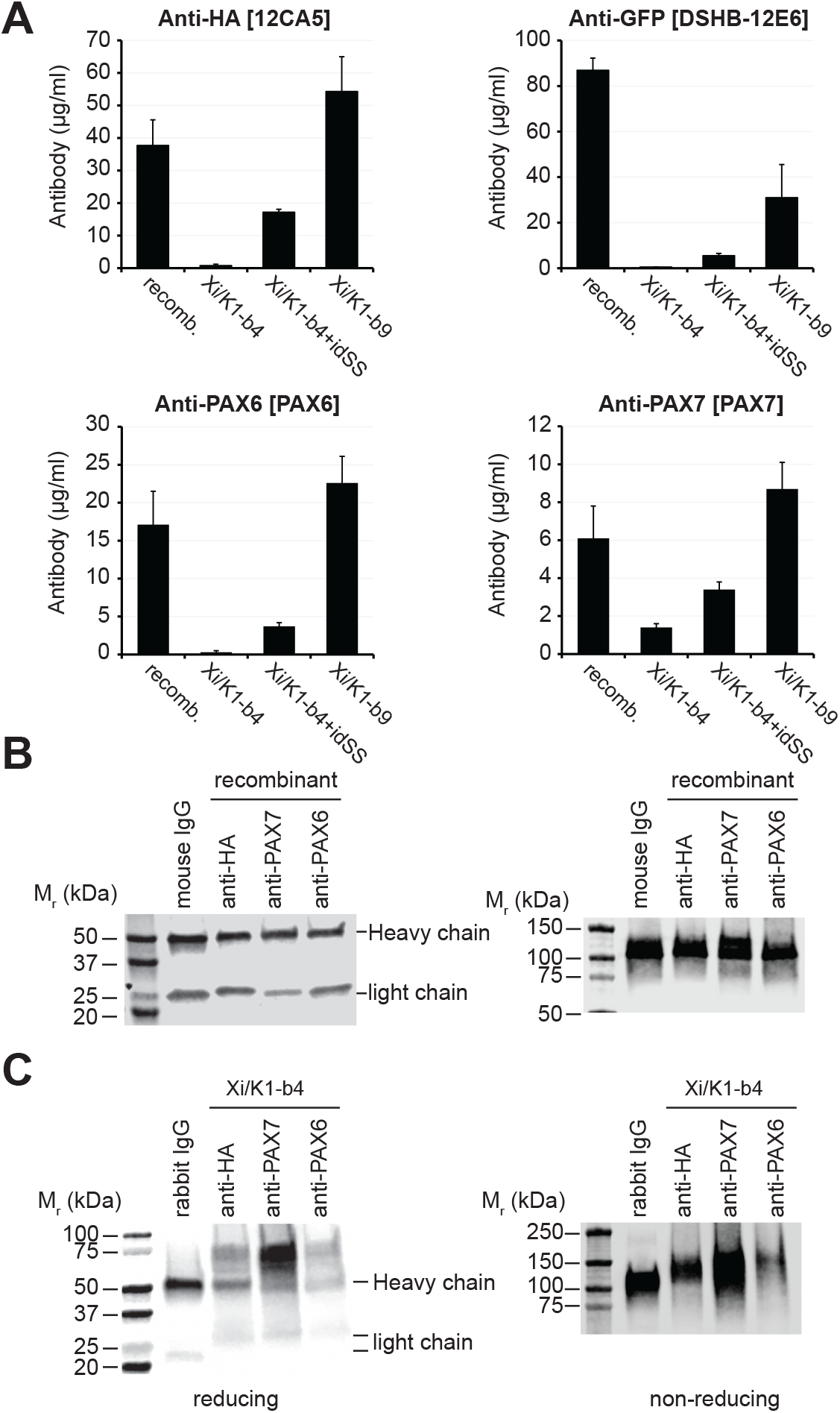
Production characteristics of chimeric and recombinant anti-bodies with engineered cysteine residues. (A) Antibody titers from native mouse recombinant antibodies (recomb.) and chimeric recombinant antibodies (Xi) using different rabbit kappa 1 (K1) light chain allotypes (b4, b9), or b4 with an interdomain disulfide bond (idSS) engineered by inserting a cysteine at position 80 in the mouse VL region (b4+idSS). Bars indicate the mean of triplicate samples; error bars represent the standard deviations. (B) Immunoblots of representative recombinant antibodies under reducing (left) and non-reducing (right) conditions. (C) Immunoblots of Xi/K1-b4 chimeric recombinant antibodies lacking the idSS under reducing (left) and non-reducing (right) conditions. Positions of the heavy and light chains are indicated; Mr, relative migration in approximate kiloDaltons (kDa). Antibodies in (B and C) were detected with anti-mouse and anti-rabbit secondaries, respectively.

These results show that chimeric recombinant mAbs using mouse variable domains heterologously in rabbit immunoglobin backbones may exhibit reduced expression levels despite robust expression as a non-chimeric mouse recombinant antibody.

### Interdomain disulfide bonds in rabbit kappa light chains are needed for robust expression of chimeric recombinant monoclonal antibodies

In rabbits, the kappa 1 light chain (K1) is the most abundant isotype and is characterized by intrachain, interdomain disulfide bonds between the variable and constant regions of the protein (**Fig. 1**; **Table 1**) (15; 25; 26). In the K1 allotypes b4, b5, and b6, disulfide bonds exist between cysteines 80 (Cys80) and 170 (Cys171), in the variable and constant regions, respectively, according to Chothia and Kabat antibody numbering (**Fig. 1**; **Table 1**). This disulfide bridge is absent in rodent and human light chains (**Table 1**), leading to a potentially unpaired cysteine 171 with a free thiol group when chimeric antibodies are made between rabbit and either human or mouse antibodies (**Fig. 1B**). This diversity in rabbit light chains has complicated the generation of chimeric rabbit-human Fabs for possible therapeutic use (27).

Because our initial chimeric recombinant antibodies were made using the more abundant K1-b4 allotype, we next determined the extent that alterations in interdomain disulfide bonds might contribute to the poor expression of mouse-rabbit chimeric antibodies. We engineered interdomain disulfide bonds (idSS) in the mouse-rabbit chimeric K1-b4 light chain by mutating amino acids at position 80 in the mouse VL (i.,e., alanine) to cysteine (**Fig. 2A**). Alternatively, we used the rabbit K1-b9 light chain allotype as the backbone for chimeric recombinant mAbs. This constant region backbone contains an alternate interdomain disulfide bridge between Cys108 and Cys171, which is unaffected by introduction of mouse variable regions (**Fig. 1**).

Reengineering a cysteine at position 80 increased the levels of the antibody production between 2-20 fold compared to those lacking this modification (**Fig. 2A**; **Table 2**), although expression levels were still below those of the respective mouse recombinant antibodies (**Fig. 2A**;**Table 2**). However, chimeric recombinant antibodies generated in the K1-b9 allotype were produced at even higher amounts, and in three of the four cases exceeded levels of their respective mouse antibodies (**Fig. 2A**; **Table 2**).

Because free thiol groups could potentially lead to aggregation or instability of recombinant antibodies (28), we next generated chimeric recombinant antibodies using rabbit variable regions in the context of the mouse light chain (rabbit-mouse chimeras; Xi/mouse), to introduce an unpaired Cys80 (**Fig. S3**). We further engineered either individual unpaired cysteines or paired Cys80 and Cys171 residues in mouse *κ* light chains (**Fig. 3A**).

**Figure 3:**
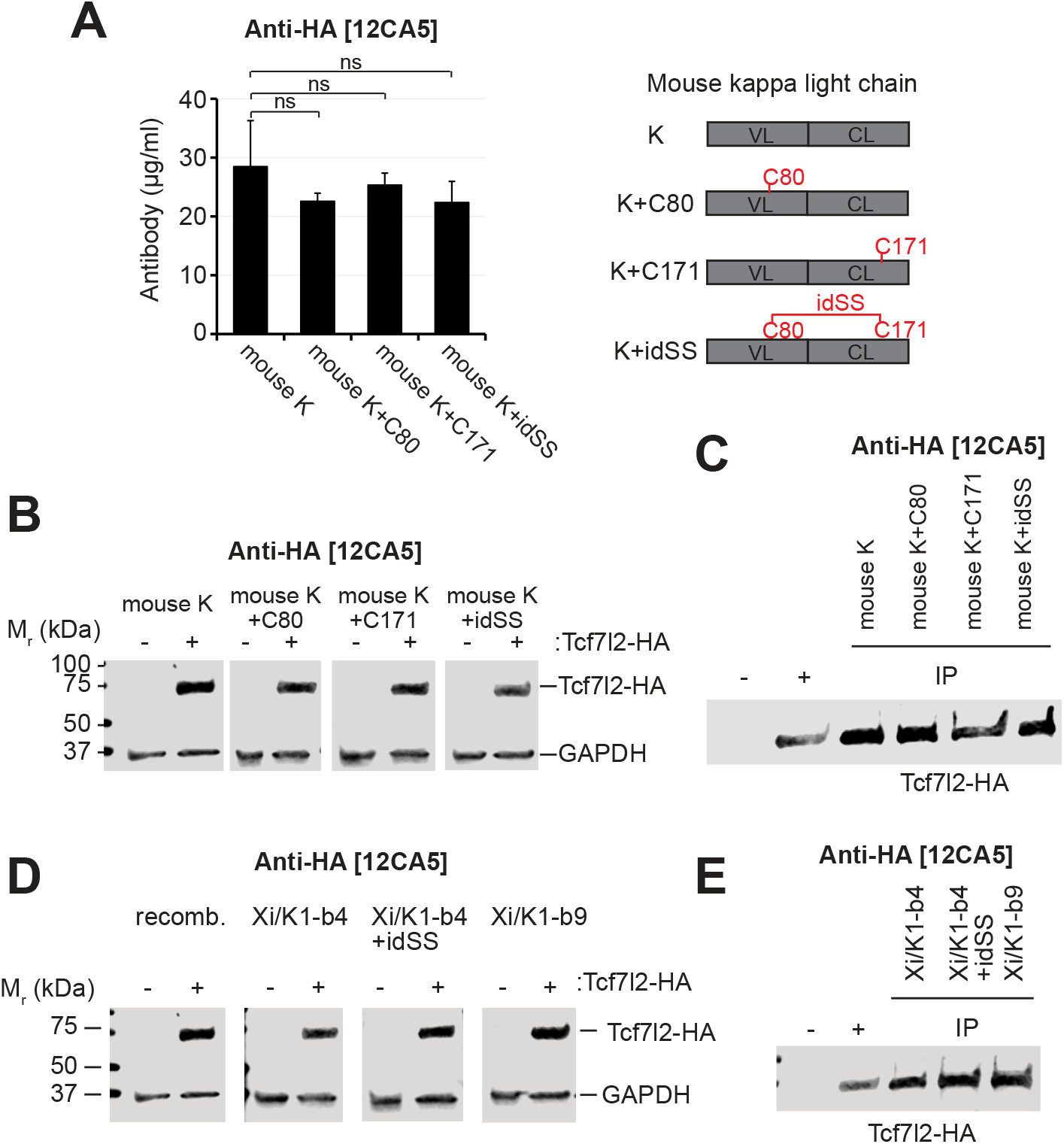
Unpaired cysteines are not sufficient to impair recombinant antibody production or activity. (A) Antibody titers from recombinant anti-HA [12CA5] mouse antibodies engineered with potentially paired or unpaired cysteines at C80 and/or C171. Bars indicate the mean of triplicate samples; error bars represent the standard deviations. n.s.=not significant (t-tests). (B) Immunoblots of engineered recombinant anti-HA [12CA5] mouse antibodies against lysates transfected with HA-tagged Tcf7l2/Tcf3 (Tcf7l2-HA). Immunoblotting of the same membrane with rabbit anti-GAPDH [2G7] was done as loading control. (C) Immunoprecipitation of tagged Tcf7l2-HA using the different anti-HA [12CA5] mAbs. (D) Immunoblots using recombinant and chimeric anti-HA [12CA5] antibodies containing different kappa light chain forms. Antibodies were used against transfected cell lysates expressing Tcf7l2-HA. Immunoblotting of the same membrane with mouse anti-GAPDH [2G7] was used as a loading control. (E) Immunoprecipitation of tagged Tcf7l2-HA using the different chimeric anti-HA [12CA5] mAbs containing different kappa light chain forms (upper panel); recombinant (recomb.) mouse anti-HA [12CA5] is shown in the lower panel.

In three different rabbit-mouse chimeras using variable regions obtained from bona fide rabbit hybridomas (CPTC-FSCN1-1, CPTC-HSPB1-6, CPTC-ANXA1-4), expression levels were similar or greater than the corresponding (recombinant) rabbit mAbs (**Fig. S3**). Similarly, introducing individual unpaired or potentially paired cysteines in the mouse light chain of Anti-HA [12CA5] did not alter expression of the recombinant antibodies (**Fig. 3A**). Furthermore, these forms were effective in recognizing overexpressed HA-tagged protein (Xenopus Tcf7l2/Tcf3-HA) in immunoblot and IP assays (**Fig. 3B-C**).

Thus, the mouse-rabbit chimeric recombinant antibodies likely require the idSS as a structural element for efficient production in human cell lines rather than for preventing free thiols per se.

### Chimeric recombinant antibody activity does not strictly depend on interdomain disulfide bridges

We next assessed the extent that the reduced production levels of chimeric antibodies might affect the binding activity or specificity towards their respective targets. For anti-HA [12CA5] mAbs, we transfected HEK293 cells with HA-tagged Xenopus Tcf3-HA and performed immunoblotting and immunoprecipitation (IP) assays. Both the original mouse anti-HA [12CA5] mAb and the different chimeric recombinant anti-HA antibodies effectively recognized the

HA-tagged protein in immunoblots of transfected samples (**Fig. 3D**). Similarly, in IP assays, recombinant and chimeric recombinant mAbs could successfully pull down the target HA-protein, roughly equivalently (**Fig. 3E**). Thus, despite the differing protein expression of chimeric mAbs, the ability to recognize target antigens remains intact for this anti-HA monoclonal antibody.

However, upon testing chimeric recombinant mAbs against PAX6 and PAX7, we found that this retention of activity was not universally preserved. We tested the activity of anti-PAX6 antibodies in immunoblotting and in IP using HEK293 cells transfected with a mouse Pax6 construct and in immunostaining against endogenous Pax6 on cryosections of chick embryos. Although all recombinant and chimeric recombinant anti-PAX6 mAbs recognized transfected Pax6 in immunoblots (**Fig. 4A**), the chimeric recombinant anti-PAX6 made using the K1-b4 allotype (K1-b4, K1-b4+idSS) detected the target with a somewhat lower signal. The K1-b4 construct failed in IP assays using transfected overexpressed Pax6 (**Fig. 4B**), whereas reintroducing a cysteine at position 80 to putatively reestablish the light interdomain disulfide bond restored IP activity. The chimeric recombinant mAb using the K1-b9 allotype were also successful in IP assays, with low levels of Pax6 present in the unbound supernatant, resembling the native hybridoma-produced IgG in the extent of Pax6 depletion from the sample (**Fig. 4B**).

**Figure 4:**
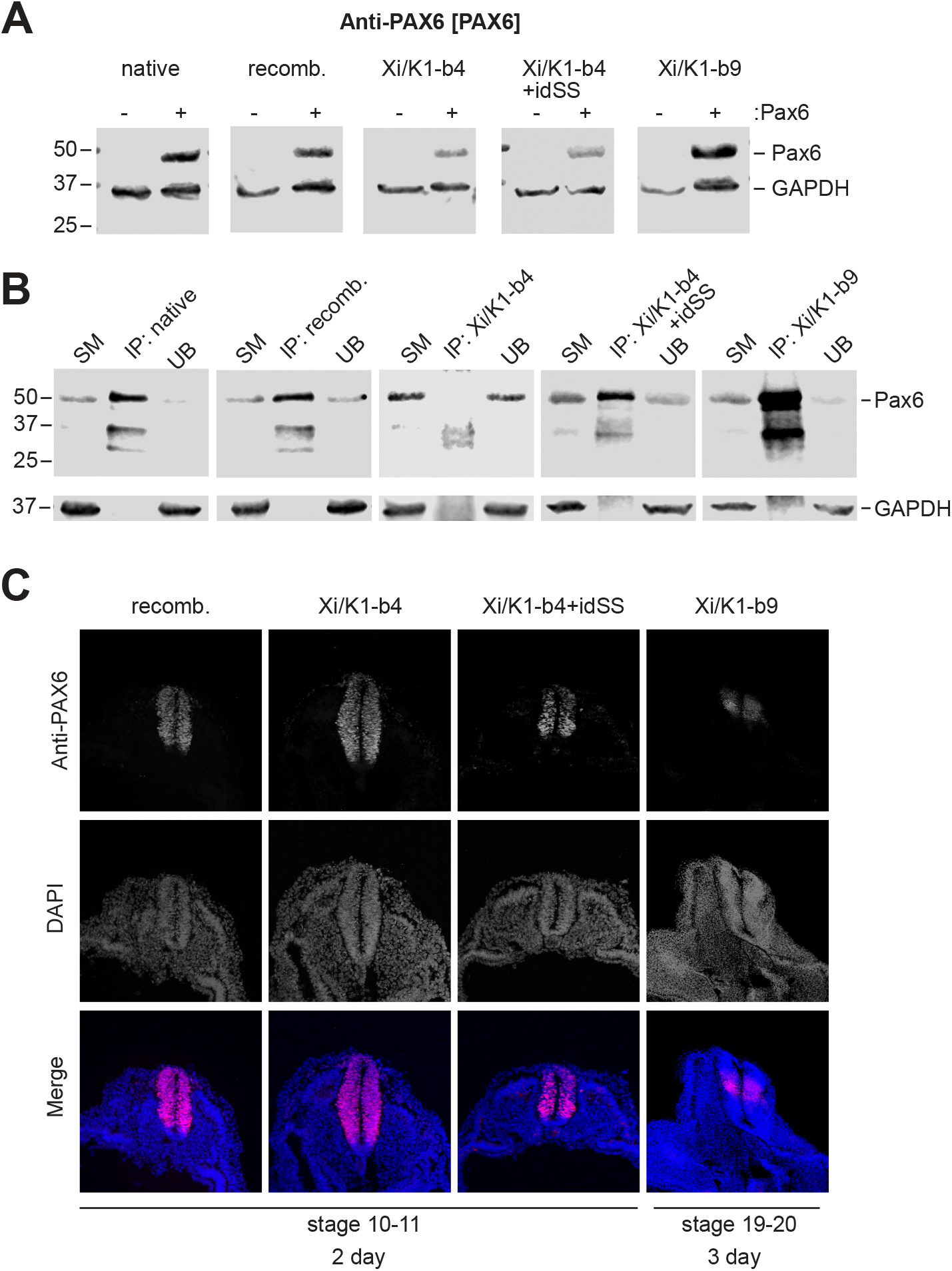
Anti-PAX6 [PAX6] chimeras retain activity in immunoblot and immunofluorescence but not for immunoprecipitation. (A) Immunoblots using recombinant and chimeric anti-PAX6 [PAX6] antibodies containing different kappa light chain forms. Antibodies were used against transfected cell lysates expressing mouse Pax6. Immunoblotting of the same membrane with rabbit anti-GAPDH [2G7] was used as a loading control. (B) Immunoprecipitation of transfected Pax6 using native, recombinant and chimeric anti-PAX6 [PAX6] mAbs. (C) Immunofluorescence against chick embryo cryosections; sections were counterstained with DAPI to visualize nuclei. Pax6 immunostaining is evident throughout the neural tube except for the ventral floor plate. Cross section view, dorsal is towards the top.

For immunofluorescence however, all recombinant and chimeric recombinant mAbs against PAX6 recognized the endogenous protein in day 2-3 chick embryo cryosections (**Fig. 4C**). In the early (stage 10) chick neural tube, PAX6 is expressed throughout the neural tube, except for the ventral most regions and floor plate (21), before becoming progressively reduced in the dorsal- and ventral-most regions in later stages (stage >18). Most notably, the staining patterns obtained using mouse recombinant mAbs and mouse-rabbit chimeric recombinant mAbs using the K1-b4 allotype were indistinguishable (**Fig. 4C**).

For anti-PAX7 antibodies, the K1-b4 chimeric recombinant mAbs (lacking Cys80) failed to recognize human PAX7 transfected into HEK293 cells in immunoblot or in IP experiments. Chimeric recombinant forms containing predicted disulfide bonds showed mostly normal immunoblotting and IP activity, with the K1-b4+idSS chimeric recombinant mAb showed detectable but greatly reduced activity in IP assays (**Fig. 5A-B**). Similarly, K1-b4 chimeric recombinant mAbs lacking Cys80 failed to detect PAX7 in chick dorsal neural tube and myotome in cryosections, whereas other forms performed similarly to the normal mouse recombinant mAb (**Fig. 5C**).

**Figure 5:**
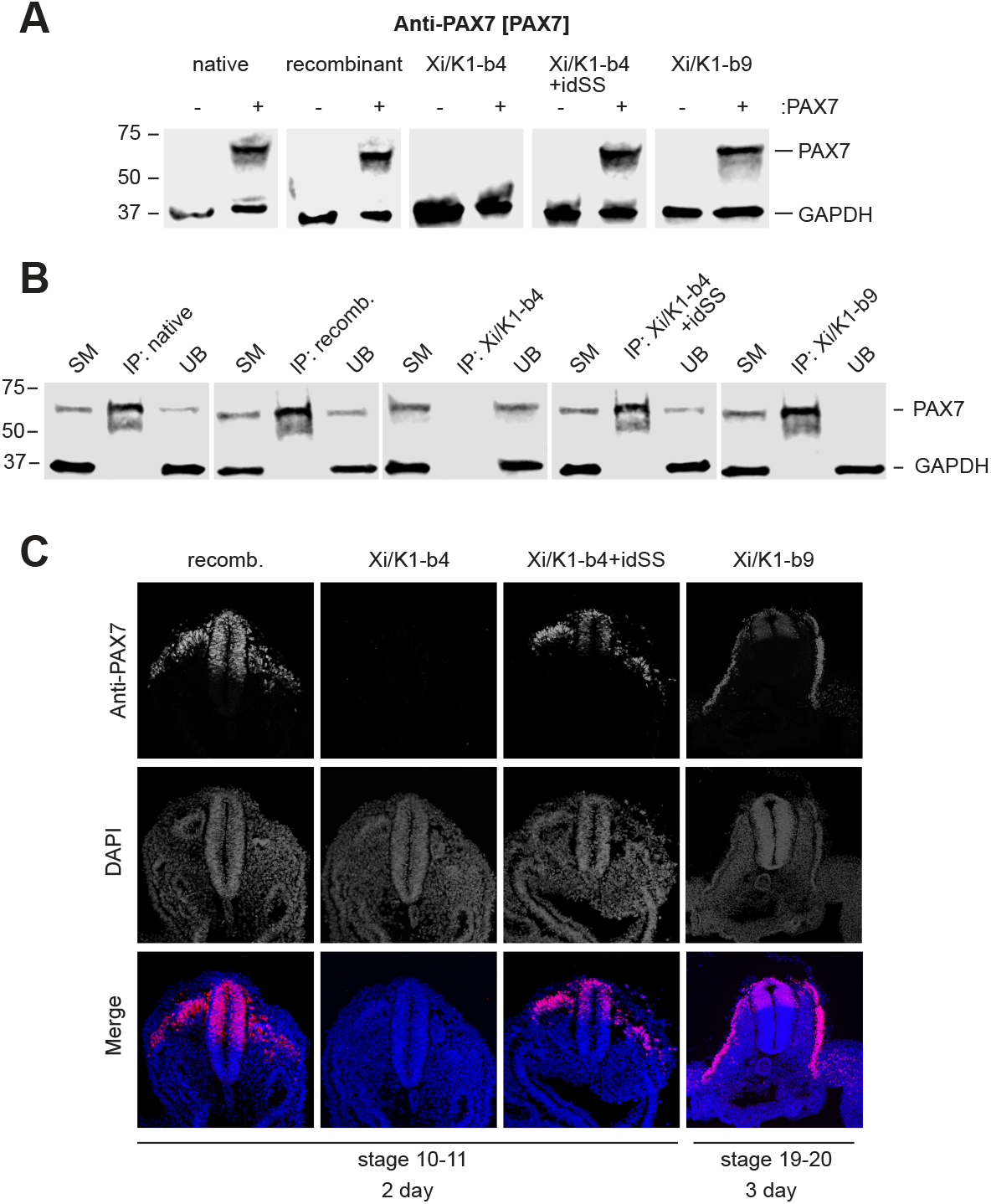
Anti-PAX7 [PAX7] K1-b4 chimeras lose activity in immunoblot, immunofluorescence, and immunoprecipitation. (A) Immunoblots using recombinant and chimeric anti-PAX7 [PAX7] antibodies containing different kappa light chain forms. Antibodies were used against transfected cell lysates expressing human PAX7. Immunoblotting of the same membrane with rabbit anti-GAPDH [2G7] used as a loading control. (B) Immunoprecipitation of transfected PAX7 using native, recombinant and chimeric anti-PAX7 [PAX7] mAbs. (C) Immunofluorescence against chick embryo cryosections; sections were counterstained with DAPI to visualize nuclei. PAX7 immunostaining is evident in the dorsal neural tube and myotome. Chimeric mAbs using the normal kappa 1 b4 allotype lack activity in all assays. Cross section view, dorsal is towards the top.

Overall, these data show that low production levels owing to the absence of the Cys80-Cys171 interdomain disulfide bond in mouse-rabbit chimeric recombinant antibodies may not necessarily predict a lack of activity in different applications, although both production levels and activity are better retained in forms containing interdomain disulfide bonds.

## Discussion

This study highlights some advantages and disadvantages of generating recombinant antibodies in antibody chains derived from different species. Overall, this work stresses the importance of an organism’s antibody repertoire in producing highly avid recombinant chimeric monoclonal antibodies while also achieving good production. Recombinant monoclonal antibodies can be made more rapidly and more cost-effectively compared to traditional hybridoma technology, and can address limitations such as clone instability and labor-intensive maintenance. Additionally, recombinant antibodies allow for a wide variety of other modifications, including isotype changes, the creation of bi/multivalent antibodies, and producing antibody fragments or single-chain antibodies. Because these recombinant antibodies can be assigned a distinct molecular identity based on amino acid sequence, there is good potential for these reagents to become true open-source resources (29), enabling better accessibility and reproducibility.

Many therapeutic and diagnostic antibodies are recombinant chimeric or humanized mouse or rabbit antibodies (30; 31). Generating chimeric mouse-rabbit and rabbit-mouse monoclonal antibodies from well-validated pre-existing antibodies is also advantageous for basic research, allowing wider applicability and potential for multiplex detection. In this study, we aimed to generate recombinant chimeric antibodies with active antigen recognition and high production yields. However, the yields of these mouse-to-rabbit chimeric antibodies (Xi/rabbit) were significantly reduced compared to their original mouse counterparts. Additionally, some recombinant chimeric antibodies showed equivalent activity to the non-chimeric antibodies in some assays (e.g., anti-HA [12CA5]), whereas others either partially (e.g., anti-PAX6) or fully (e.g., anti-PAX7) lost the ability to bind the target protein. Other reports have noted similar findings for mouse-rabbit recombinant chimeric antibodies (32), but the underlying causes remain unclear.

We examined mouse VL regions and the rabbit CL regions for potential incompatibilities between antibodies from these species. Our initial chimeras used the rabbit K1-b4 allotype, the most abundant light chain form in the immunoglobin repertoire of New Zealand white rabbits. The b4 allotype, along with the b5 and b6 allotypes, is characterized by the presence of an interdomain disulfide bond between VL (Cys80) and CL (Cys171). This feature is not found in human or mouse antibodies. It has been suggested that this bond may enhance the stability and antigen affinity of rabbit Fab molecules (18; 33), although one study showed that removing this disulfide bond in rabbit antibodies only slightly reduces thermal stability, and had no apparent impact on their antigen affinity (19).

Our work in this study showed a dramatic decrease in the production of chimeric recombinant antibodies that lack the cysteine at position 80, absent from the mouse VL, and thus likely lack the Cys80-Cys171 interdomain disulfide bond. Reintroduction of a Cys80 in the mouse VL to presumptively restore this interdomain disulfide bond increased production by between 2-20 fold. Despite this improvement, expression levels of these chimeric antibodies remained lower than those of their respective recombinant mouse antibodies, indicating that other mechanisms may be involved as well. Unique among rabbit light chains, the K1-b9 allotype has an interdomain disulfide bond between Cys108, the first amino acid after the VL domain, and Cys171 in CL (**Figs.1, 2**; **Table 1**), and is not affected by replacement of the rabbit VL by that of mouse. Thus, using the K1-b9 allotype in recombinant chimeric antibodies eliminates the need for re-engineering cysteine modifications to achieve sufficient expression levels in transfected cells.

Additionally, the activity of the anti-PAX6 and anti-PAX7 chimeric antibodies was impaired under some conditions. Notably, introduction of the interdomain disulfide bond into the chimeric K1-b4 light chains of these chimeric antibodies increased production from 2 to 20 fold. Despite this improvement, expression levels of these chimeric antibodies remained lower than those of their respective recombinant mouse antibodies, indicating that other parameters may need optimization as well. Use of the K1-b9 allotype eliminates the need for cysteine modification at position 80 for idSS formation, and these K1-b9 chimeric antibodies achieved expression levels similar to those of their respective mouse antibodies. Additionally, the interdomain disulfide bond restoration improved antigen recognition in immunoblotting, IP, and immunostaining. These findings underscore the importance of the interdomain disulfide bond in enhancing chimeric antibody production and activity. Furthermore, comparing the alterations in sequence and/or structure may be useful in prediction of antigen recognition by different antibody sequences using artificial intelligence tools.

Many therapeutic humanized antibodies have been developed using CDR grafting (34; 35). Applying this technique to graft CDRs into the framework of rabbit antibodies may avoid the problems of antibody stability and lower production levels, but this method is more technically challenging and requires a more complete knowledge of the key recognition residues. We ruled out any negative impacts of unpaired cysteine residues generated in chimeric antibodies on production and antigen affinity. And, we confirmed that introducing an extra uncoupled cysteine, either C80 or C171, into mouse light chain antibodies did not significantly affect their production or activity. These observations suggest that the idSSs are important structurally, and not merely to prevent free thiol groups.

Similar to variation in antibody levels produced by traditional hybridomas, we saw considerable variability in production levels among recombinant antibodies generated in this study. This variability could be attributed to differences in the signal peptides used for expressing each antibody. Therefore, swapping the signal peptides, which have a significant influence on antibody secretion into the growth medium (36), may enhance antibody production. Recombinant antibody yields may also be influenced by other factors, such as the choice of cell lines, cell fitness conditions, and transfection reagents, as well as their primary sequences deciding molecular stability. Chimeric K1-b4 antibodies produced low amounts of intact antibody in cell culture supernatants, and showed gradual aggregation after being concentrated prior to IP applications. Antibody aggregation could indicate aberrant folding or instability, revealing hydrophobic cores prone to aggregation (37; 38). Thus, even chimeras that effectively retain antigen recognition may have limited use owing to the behavior of the secreted molecules.

Rabbit monoclonal antibodies offer several advantages including high affinity and specificity over mouse antibodies (39), and tend to recognize diverse immunogens including those poorly or non-immunogenic in mice and humans. In this study, we show that rabbit variable regions can be cloned into mouse immunoglobulin backbones without the need for additional modifications to the primary sequences, whereas this is not the case for the reverse direction. These results suggest that various recombinant antibody variants from rabbit monoclonal antibodies can be easily engineered and produced, once the sequences of variable regions of rabbit monoclonal antibodies are obtained through various methods, such as phage-display, single B-cell sequencing, NGS techniques, and sequencing of cDNA from rabbit hybridomas (11; 40). Because bona fide rabbit hybridomas tend to exhibit instability and thus lose antibody secretion over time (41; 42), recombinant antibody engineering should prove to be exceptionally valuable in preserving these antibodies.

In summary, our study highlights the importance of taking into account species-specific antibody features when engineering chimeric antibodies. Similar considerations could be applied to antibody engineering across various species.

## Materials and methods

### Antibodies

Antibody anti-HA [12CA5] (RRID: AB_3105930) was obtained from DSHB at the University of Iowa, originally deposited by DSHB in 2022. Antibody anti-GFP [DSHB-12E6] (RRID: AB_2617418) was obtained from DSHB at the University of Iowa, originally deposited by DSHB in 2013. Antibody anti-PAX6 [PAX6] (RRID: AB_528427) was obtained from DSHB at the University of Iowa, originally deposited by Kawakami, A. in 1997. Antibody anti-PAX7 [PAX7] (RRID: AB_528428) was obtained from DSHB at the University of Iowa, originally deposited by Kawakami, A. in 1997. Antibody anti-GAPDH [DSHB-2G7] (RRID: AB_2617426) was obtained from DSHB at the University of Iowa, originally deposited by DSHB in 2016. Antibody CPTC-FSCN1-1 (RRID: AB_1553796) was obtained from DSHB at the University of Iowa, originally deposited by the Clinical Proteomics Technologies for Cancer program in 2020. Antibody CPTC-HSPB1-6 (RRID: AB_2617271) was obtained from DSHB at the University of Iowa, originally deposited the Clinical Proteomics Technologies for Cancer program in 2014. Antibody CPTC-ANAX-4 (RRID: AB_2617223) was obtained from DSHB at the University of Iowa, originally deposited the Clinical Proteomics Technologies for Cancer program in 2014.

### RNA isolation and sequencing of hybridoma VL and VH domains

Total RNA was isolated from approximately 1 × 10^7^ actively growing hybridoma cells using the RNeasy kit (QIAGEN, Catalog*#* 74004) following the instructions provided by the manufacturer. To synthesize cDNAs corresponding to antibody variable heavy (VH) and variable light (VL) chain regions, reverse transcription-polymerase chain reaction (RT-PCR) was performed using the SMARTScribe kit with chain specific primers (**Table S1**), according manufacturer’s instructions (Takara Bio, Catalog# 639537) and methods in (43). Conventional PCR was then used to amplify VH and VL regions. The resulting blunt end PCR products were cloned into the pCR-XL-2-TOPO vector (Invitrogen, Catalog# K8050). The cloned PCR products were subsequently sequenced using Sanger sequencing. The identical sequences were verified in at least three clones to ensure accuracy.

### Cloning of recombinant and chimeric recombinant antibodies

The pcDNA3.4 vector, a constitutive mammalian expression vector, was used to construct separate expression plasmids for the heavy and light chains of each antibody. VH and VL chain fragments were PCR amplified using antibody-specific primer pairs (**Table S1**) from primary pTOPO vectors, which contained VH and VL sequences derived either from cDNA or DNA gBlock™ fragments synthesized by Integrated DNA Technologies, Inc. (IDT, Coralville, IA). The constant regions of the mouse IgG1 heavy and kappa light chains, as well as the rabbit IgG heavy and kappa 1-b4 and -b9 light chains, were synthesized as DNA gBlocks (IDT) and cloned into the pCR Blunt II-TOPO vector (Thermo Fisher, Catalog# K280002). The DNA sequences for the constant regions of mouse and rabbit gamma and kappa immunoglobulins were obtained from NCBI with following accession numbers: AAB59656 (mouse IgG1 heavy chain), Z27396 (mouse kappa light chain), XP_069921742 (rabbit IgG heavy chain), X00231 (rabbit kappa1-b4 light chain allotype), K01359 (rabbit kappa1-b9 light chain allotype). The constant heavy (CH) and constant light (CL) fragments were PCR amplified from the respective pTOPO vectors using primers listed in **Table S1**, incorporating appropriate restriction sites. The resulting variable and constant fragments were digested with restriction enzymes and ligated into the pcDNA3.4 vector.

To introduce cysteine modifications at position 80 and 171 in the light chains, site-direct mutagenesis was performed using the PCR-driven overlap extension method (44).

All amplified sequences in the expression plasmids were verified through Sanger sequencing. Large-scale preps of plasmid DNA were prepared and the quality assessed prior to transfection experiments.

### Recombinant antibody production

Two cell lines, HEK293C18 (ATCC; CRL-10852, RRID:CVCL_6974) and Expi293F (Thermofisher, Catalog# A14635), were used for transient transfection and in vitro expression of recombinant antibodies. For both cell lines, plasmids encoding the heavy and light chains for each antibody were mixed at a 1:2 ratio and transfected using appropriate reagents.

For transfection of HEK293C18 cells, the plasmids were transfected using PEI (Polyethylenimine) (Polysciences, Catalog# 9002-98-6, 26913-06-4). The cells were grown in a suspension culture consisting of a 1:1 mixture of FreeStyle 293 expression medium (Gibco) and SFM4HEK293 medium (HyClone) supplemented with 50 µg/ml geneticin. The transfection procedure with PEI as described in previous studies (45; 46), modified and optimized for antibody production.

For Expi293F cells, transfection was performed using Expi293™ Expression System Kit (Thermofisher, Catalog# A14635), following the manufacturer’s protocol for both cell culture and transfection.

After seven days incubation, the supernatant containing the expressed antibodies was harvested and the antibody concentration was quantified by ELISA.

### Recombinant protein expression

Recombinant proteins were expressed in HEK293C18 cells using transient transfection with PEI (see previously). Transfected cells were collected after two days of incubation and lysed in a lysis buffer containing protease inhibitors. The resulting cell lysates were then subjected to further immunoblotting and immunoprecipitation analysis. Plasmids for transfection were Xenopus Tcf7l2/XTcf3 tagged with HA at the C-terminus (and FLAG at the N-terminus), mouse Pax6, and human PAX7 tagged with FLAG at the N-terminus. pCS2+Flag-XTCF3-HA was a gift from Sergei Sokol (Addgene plasmid # 32998; http://n2t.net/addgene:32998; RRID:Addgene_32998). pcDNA3.4-Pax6 was generated in this study. Mouse Pax6 was amplified from the plasmid pMXs-Pax6, a gift from Kevin Eggan (Addgene plasmid # 32932 ; http://n2t.net/addgene:32932; RRID:Addgene_32932). FLAG-hPAX7 (#348) was a gift from Matthew Alexander & Louis Kunkel (Addgene plasmid # 78337; http://n2t.net/addgene:78337; RRID:Addgene_78337) (47; 48).

### Immunoblotting

Immunoblots were performed as previously described (49; 50). Briefly, denatured protein samples or reduced/non-reduced antibodies were separated on 4-15% SDS-PAGE gels (Bio-Rad). Subsequently, the separated proteins were blotted onto a 0.22 µm nitrocellulose membrane (Amersham). The membranes were blocked in a 5% non-fat milk solution, and then incubated with primary antibodies at a concentration of 0.5 µg/ml in a blocking buffer. Blotting against GAPDH was used to control for equal loading, using either mouse or rabbit anti-GAPDH mAbs. Incubation was carried out either for 2 hours at room temperature or overnight at 4°C with rocking. After three washes with TBS-T (TBS + 0.05% Tween-20), the membranes were incubated with IRDye680/800-conjugated goat anti-mouse and/or goat anti-rabbit secondary antibodies (Li-Cor Biosciences) at a 1:10,000 dilution for 1 hour at room temperature. The membranes were washed with TBS-T and scanned using an Odyssey Fc Dual Mode Imaging System (Li-COR Biosciences).

### Immunoprecipitation

Immunoprecipitation (IP) was performed as previously described (49). Transfected cells (about 1 × 10^8^) were pelleted and lysed in 1000 µl of lysis buffer (Cell Signaling Technology, #9803) with protease inhibitor cocktail (Roche) for 15 minutes on ice. Cell debris was removed by centrifugation at 10,000 rpm (9,600 x g) for 8 minutes. Total protein concentration in the clarified lysates was determined using the Pierce BCA Protein Assay Kit (Thermofisher, Catalog# A55865).

For immunoprecipitation of the target antigen, diluted lysates (∼ 1 mg total protein) were incubated with 5-8 µg of antibody pre-bound to protein A/G magnetic beads (500 µg, Bimake.com) for 30 minutes at room temperature or for 1 hour at 4°C with rotation. After incubation, aliquots of input (the starting material) and unbound fractions were kept for later analysis. Beads were washed with wash buffer (PBS + 0.01% Tween-20). The eluates were separated by SDS-PAGE and analyzed by immunoblotting.

### Immunostaining

Fertilized chicken eggs were incubated at 38°C for 2-3 days. Immunostaining was performed using methods modified from (51). Briefly, embryos were harvested and fixed for 2 hours in 4% paraformaldehyde in 0.1 M phosphate buffer, pH 7.2. After washing in ice-cold PBS, the embryos were transferred to cold 30% sucrose in 0.1M phosphate buffer and allowed to equilibrate overnight. Each embryo was embedded with tissue freezing medium (OCT) in a silicon mold and frozen on dry ice and then transferred to −80°C freezer. Frozen embryos were sectioned to a thickness of 20 µm, mounted on charged Superfrost plus microscope slides (Fisher Scientific), and allowed to dry at room temperature. Sections on slides were rehydrated in PBS for 15 min and then blocked in PBS-T (1.0% goat serum, 0.1-1% Triton X-100 in PBS) for 1 hour at room temperature. Incubation with primary antibodies diluted to a concentration 1 µg/ml in PBS-T was performed in a humidity chamber overnight at 4°C. After three washes for 5 min in PBS-T, sections were incubated with anti-mouse or anti-rabbit secondary antibodies conjugated with Cy3 (Jackson ImmunoResearch, Catalog #s 115-165-166, 111-165-144) at 1:500 dilution and 1 µg/ml DAPI. After three washes, sections were mounted in an antifade mounting medium (Vector laboratories) and stored in a dark chamber at 4°C until imaging.

### Imaging

All immunostaining images were acquired using a Leica TCS SP8 inverted confocal microscope incorporated into a Leica imaging system with LAS X software. Z-series images were acquired at intervals of 0.6 µm using a 20x objective. DAPI and Cy3 fluorescence markers were excited using laser lines at 405 nm and 552 nm respectively. The obtained *\**.*lif* file z-series images for each channel were subjected to z-projections and subsequently converted into TIFF image format using FIJI software (52). Images for publication were prepared in Adobe Illustrator and Photoshop using only level and contrast adjustments applied over the entire image. Other modifications included resizing, changing stroke/fill weights and colors and annotated overlays.

## Data availability

The data supporting the findings of this study are available within the article and/or its supporting information. The antibodies described in this study are deposited with the Developmental Studies Hybridoma Bank (DSHB; https://dshb.biology.uiowa.edu/).

## Acknowledgments

We thank DSHB team members for critical feedback and for expert technical advice on antibody production.

This work was funded in part by a grant from the NIH/ORIP, R24OD037757 to D.W.H. The content is solely the responsibility of the authors and does not necessarily represent the official views of the National Institutes of Health.

## Author contributions

Y.-N. Park: Methodology, Visualization, Writing – initial draft

D.W.H: Conceptualization, Funding acquisition, Methodology, Project administration, Supervision, Visualization, Writing – review & editing

## Supplementary material

**Figure S1:**
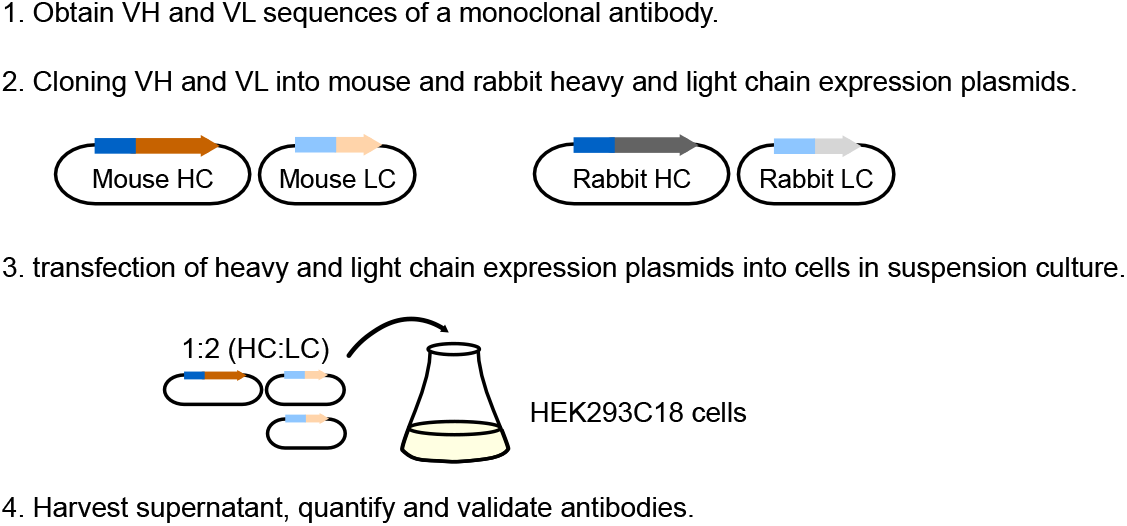
Overview of recombinant antibody production procedure

**Figure S2:**
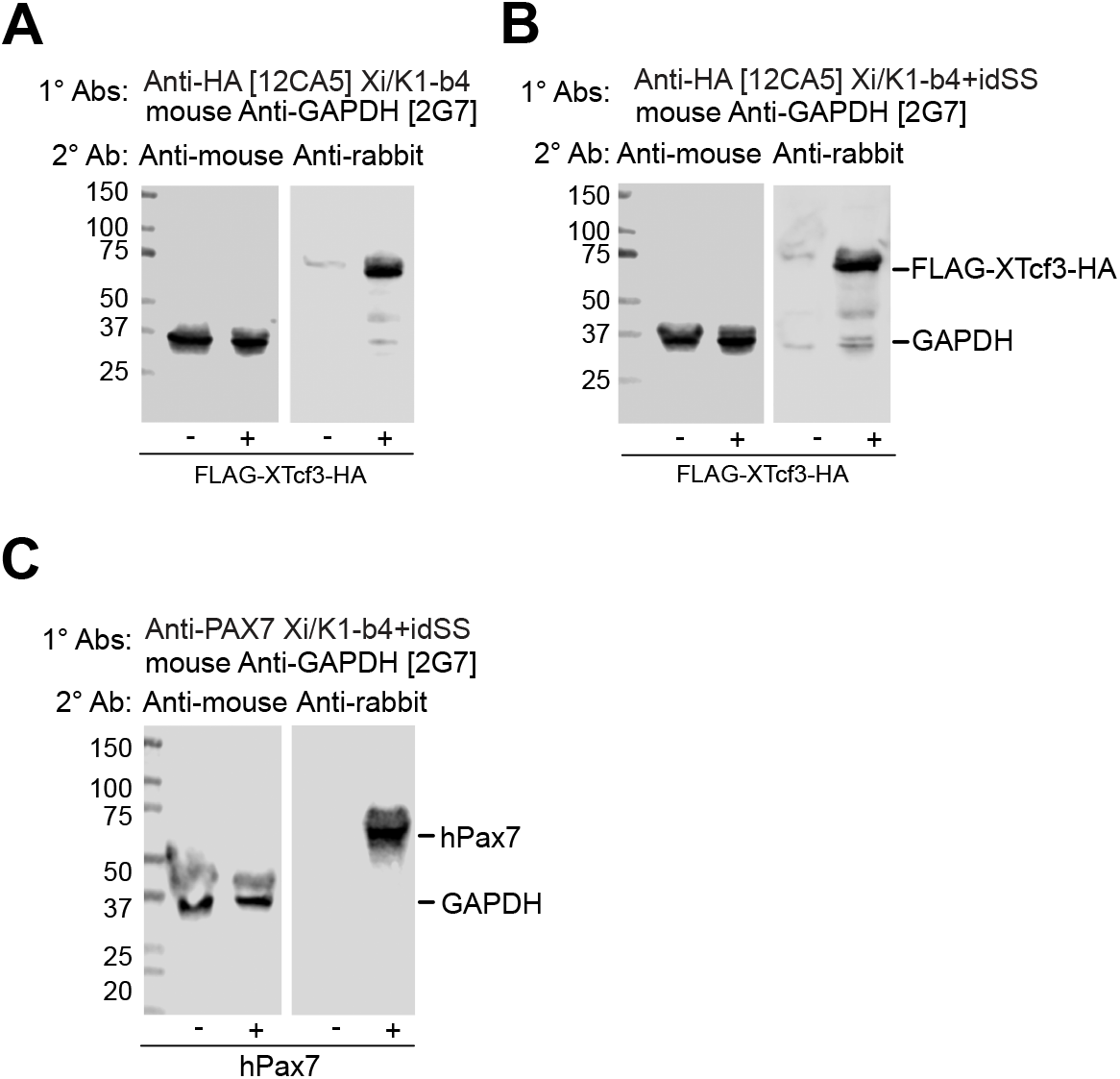
Anti-mouse secondary antibodies do not detect mouse variable domain sequences in chimeric antibodies. (A-B) Immunoblots of cells transfected with Tcf712-HA (FLAG-XTcf3-HA) using K1-b4 chimeric mAbs (A) or the same chimera with the potential idSS (B). Mouse anti-GAPDH [2G7] was used as a loading control and to show activity of the mouse secondaries. (C) Immunoblots of cells transfected with human PAX7 using the K1-b4+idSS chimera. Mouse anti-GAPDH [2G7] was used as a loading control and to show activity of the mouse secondaries.

**Figure S3:**
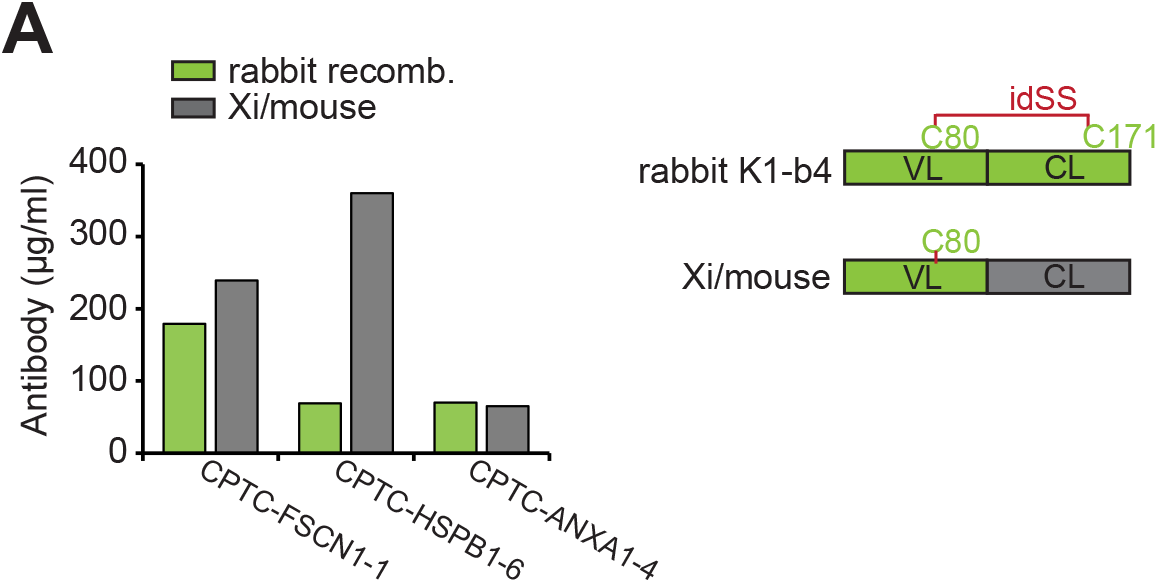
Unpaired cysteine 80 in rabbit monoclonal recombinant antibodies do not impair recombinant antibody production or activity. (A) Antibody titers from native rabbit recombinant antibodies (derived from rabbit hybridomas) and chimeric recombinant antibodies (Xi/mouse). A diagram of the respective light chains is shown on the right, showing the putative unpaired C80.

**Table S1:**
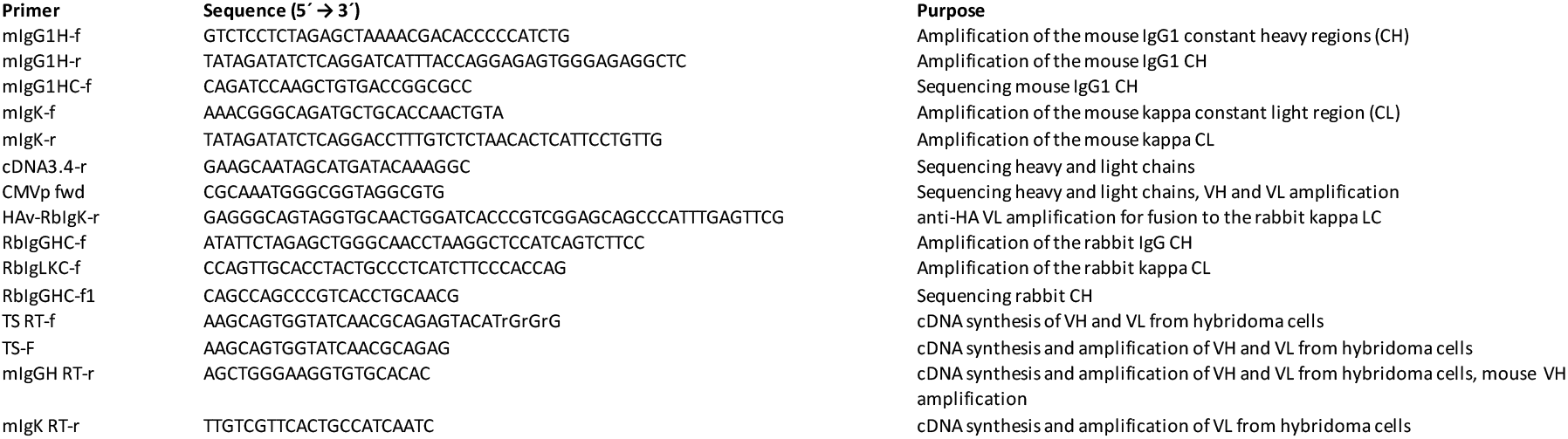
Primers used in this study.

